# PI3K/mTOR activity sensitizes cancer cells to nucleolar stress

**DOI:** 10.1101/2025.11.12.688066

**Authors:** Myriam Barz, Alba Corman, Maria Häggblad, Alicia Gonzalez-Serrano, Louise Lidemalm, Matilde Murga, Daniela Huhn, Oscar Fernandez-Capetillo

## Abstract

Drugs targeting nucleoli and generating nucleolar stress (NS) such as RNA Polymerase I inhibitors have shown anticancer properties with some progressing to clinical trials. However, the mechanisms that modulate the sensitivity to NS remain poorly understood, which has limited the design of clinical trials on patients most likely responding to these therapies. To get a panoramic view of the genetic determinants that shape the response to NS in cancer cells, we conducted genome-wide CRISPR screens in cells treated with 2 independent RNA Polymerase I inhibitors: Actinomycin D (ActD) and BMH-21. As expected, mutations known to regulate the response to NS such as *P53* or *RB1* increased the resistance to both drugs. On the other hand, the toxicity of RNA Pol I inhibitors was increased in the context of mutations that enhance PI3K/mTOR signaling. Surprisingly, we found mutations that sensitized to ActD but increased the resistance to BMH-21, revealing that, while both drugs are used as RNA Pol I inhibitors, the must have additional unknown targets. Together our study provides a global landscape of the mechanisms that modulate the sensitivity to NS-inducing agents and illustrate the need for an in-depth analysis of the mechanism of toxicity of drugs, particularly when these are advancing to clinical use.

## INTRODUCTION

The nucleolus plays a central role in the regulation of cellular homeostasis, primarily by coordinating ribosomal RNA (rRNA) transcription and ribosome assembly(Gonzalez-Arzola, 2024; Munoz-Velasco *et al*, 2025). As a result, the nucleolus is highly sensitive to various forms of cellular stress, including those that disrupt ribosome biogenesis, a condition termed *nucleolar stress* (NS) (Boulon *et al*, 2010; James *et al*, 2014; Yang *et al*, 2018). NS triggers a signaling cascade that leads to cell cycle arrest, apoptosis, or senescence, largely through the stabilization of tumor suppressor proteins such as p53 (Rubbi & Milner, 2003). The disruption of nucleolar function has therefore emerged as a promising therapeutic avenue in oncology, particularly since cancer cells often exhibit elevated ribosome biogenesis to sustain their high proliferation rates (Bhat *et al*, 2015; Carotenuto *et al*, 2019; Corman *et al*, 2023).

Among the strategies to target nucleoli in oncology, those targeting RNA Polymerase I (Pol I), the enzyme responsible for rRNA transcription, have attracted a particular interest. Inhibitors of Pol I, such as Actinomycin D (ActD) and the more recently developed BMH-21, disrupt nucleolar homeostasis and impair cancer cell viability (Chen *et al*, 2019; Drygin *et al*, 2009; Ferreira *et al*, 2020; Mars *et al*, 2020). ActD intercalates DNA and inhibits transcription broadly but at low doses selectively suppresses rRNA synthesis, while BMH-21 directly induces degradation of the Pol I complex without DNA intercalation. Some of these inhibitors have progressed into clinical trials. However, despite the emerging interest in this field, the lack of comprehensive understanding of the genetic determinants that modulate the response to NS-inducing agents has limited the capacity to stratify patients on the basis of who would most likely benefit from these treatments.

To address this gap, we conducted genome-wide CRISPR-based loss-of-function screens in human cells treated with either ActD or BMH-21. By using two mechanistically distinct Pol I inhibitors, we aimed to obtain an unbiased map of genetic dependencies and resistance mechanisms linked to NS. Our findings not only confirmed previously established modulators such as *TP53* and *RB1*, but also identified novel gene-drug interactions, including a higher sensitivity to NS in the context of mutations that increase PI3K/mTOR pathway activation. Interestingly, certain genetic perturbations conferred opposite sensitivities to ActD and BMH-21, indicating that, while both drugs induce NS, the response to these agents differs substantially. Collectively, our study provides a functional genomic landscape of the response to NS and suggests potential biomarkers that could be of use for the clinical development of RNA Pol I inhibitors in oncology.

## RESULTS

### A genome-wide screen to identify modulators of the NS-response

To obtain a panoramic view of the genes and pathways that modulate the response to NS-inducing agents, we performed a genome-wide CRISPR screen in A375 human melanoma cells. A375 cells expressing Cas9-BFP were independently transduced twice with the Brunello library, which targets ∼20,000 genes with 4 sgRNAs per gene in two independent biological replicates (Doench *et al*, 2016) (**Fig. S1A**). After puromycin selection, cells were harvested and seeded in media containing either DMSO (control), ActD (0.75 nM), or BMH-21 (0.2 μM) as 2 independent agents inducing NS (**Fig. 1A, B**). To ensure 1,000-fold library coverage, 80 million cells were seeded per condition, and this coverage was maintained through successive harvesting and reseeding into fresh compound media every three days for up to 12 days. This design allowed both longitudinal sampling and sustained selective pressure. Early timepoints were represented by samples reaching ∼50% viability (IC50) under each treatment—day 3 for ActD and day 6 for BMH-21—capturing genes whose knockout sensitizes cells to NS (**Fig. 1C**). While samples harvested on the late timepoint day 12 captured mutations providing longer-term resistance or adaptation.

**Figure 1.**
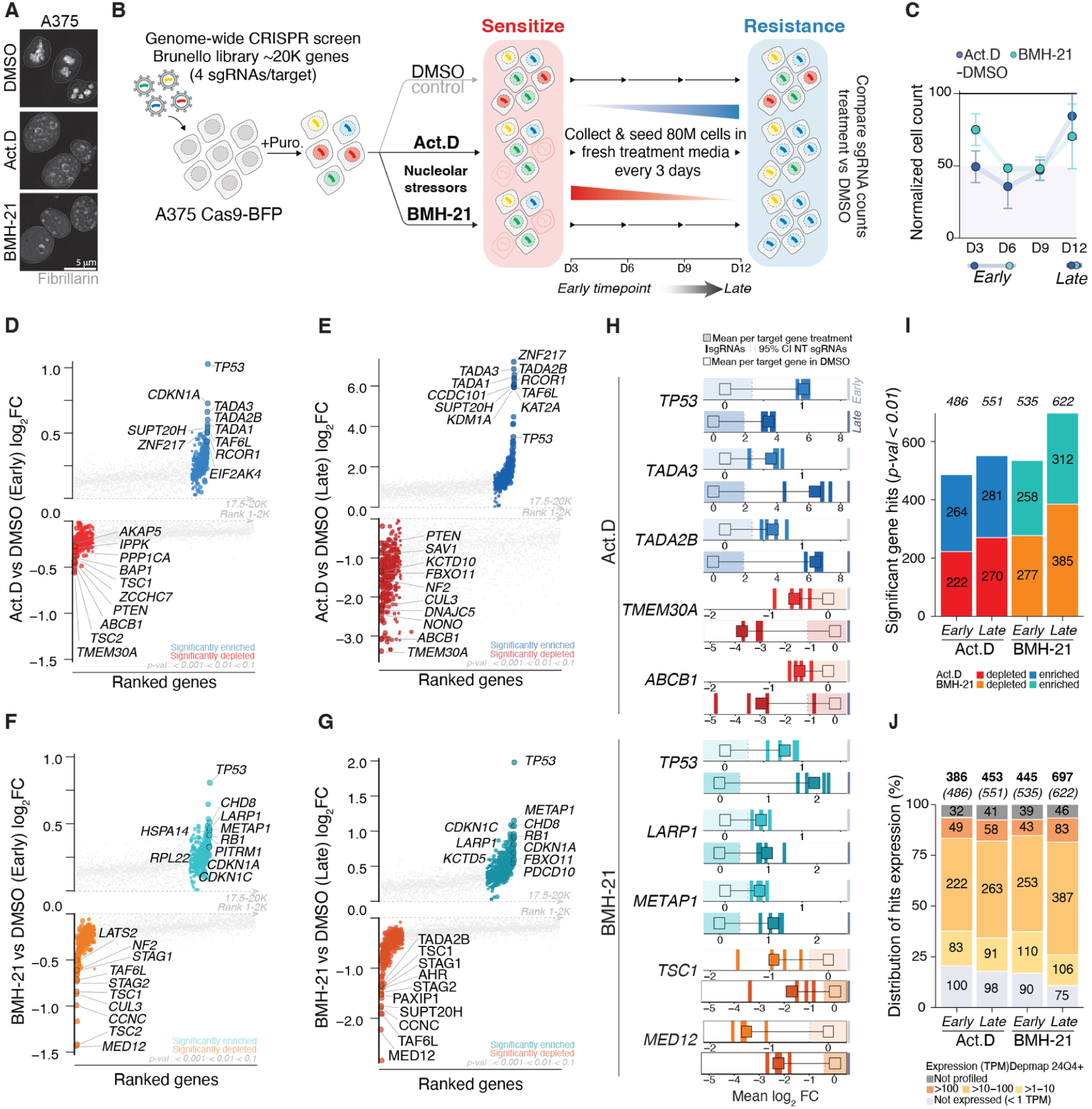
Genome-wide CRISPR screens of ActD and BMH-21 sensitivity. (A) Immunofluorescence of the nucleolar marker FIBRILLARIN. A375 cells were treated with 5 nM ActD or 2 μM BMH-21 for 6 hr. FIBRILLARIN is shown in grey; nuclei were counterstained with DAPI. Scale bars, 5 μm. (B) Schematic of the nucleolar stress genome-wide CRISPR screen workflow. (C) Normalized cell counts relative to DMSO-treated in A375 cells treated with ActD or BMH-21. At later timepoints, partial recovery of cell numbers likely reflects the emergence of resistant populations. (D–G) Plots representing log2 fold-change (log2FC) in sgRNA abundance for genes conferring resistance (enriched) or sensitivity (depleted) compared to DMSO controls at early and late timepoints for ActD (blue = enriched, red = depleted) and BMH-21 (teal = enriched, orange = depleted). To improve clarity, only the top 2,000 depleted and bottom 2,500 enriched ranks are shown. Dot size reflects RRA p-value significance, and the top/bottom 10 most significant hits are annotated for each condition and timepoint. (H) Fold-change (log2FC) values for all sgRNAs targeting each gene are shown for two independent biological replicates. Striped bars denote the number of sgRNAs per target. Filled colored squares show the mean log2FC per target gene under treatment; unfilled squares indicate the DMSO baseline (log2FC ≈ 0). A shaded area shows the 95% confidence interval for non-targeting (NT) sgRNAs under each treatment condition, highlighting the distribution of neutral controls. (I). Bar plots illustrating the number of significant hits (RRA p < 0.01) for each treatment and timepoint, as well as the total per condition. (J) Distribution of significant hits across expression tiers (not expressed <1 TPM; low 1–10 TPM; moderate 10–100 TPM; high >100 TPM) using batch-corrected DepMap 24Q4 expression data. For each condition, the final number of expressed hits (TPM > 1) is shown in bold above each bar, with the original total number of significant hits shown below.

Quality controls confirmed the robustness of the screen. The distribution of sgRNAs in the plasmid library demonstrated genome-wide representation of targets, and correlations among replicates indicated high consistency between biological duplicates. Moreover, comparison of counts from cells carrying targeting versus non-targeting (NT) guides revealed clear phenotypic divergence, more pronounced under ActD than BMH-21 (**Fig. S1B-D**).

Gene-level scores were computed using robust-rank aggregation (RRA) in MaGeCK (Li *et al*, 2014) based on guide enrichments relative to NT controls under each treatment (ActD or BMH-21) and then compared to DMSO. This analysis identified mutations conferring either resistance or sensitivity to ActD and BMH-21 at early and late timepoints (**Fig. 1D-G, Table S1**.). Consistent with its known role in the NS response, *TP53* deletions emerged as a resistance hit across all conditions. Loss of the *ABCB1* multidrug-resistance pump sensitized cells to ActD, as expected since ActD is an ABCB1 substrate (Hill *et al*, 2013). These analyses identified other interesting aspects such as the fact that the members of the SAGA complex (e.g. *SUPT20H, TADA3, TADA2B, etc.*) conferred resistance to ActD but sensitivity to BMH-21. Interestingly, depletion of negative regulators of PI3K/mTOR signaling (e.g. *PTEN, TSC1, TSC2*) increased sensitivity to both agents, while activation of the pathway (*LARP1*) reduced its toxicity. These effects were consistent across sgRNAs and became more evident over time (**Fig. 1H, Fig S1E**). Furthermore, this analysis revealed that most hits were non-essential, as they remained within the 95% confidence interval of non-targeting controls in DMSO (**Fig. 1H, Fig S1E**). On average, the screen identified approximately 500 significant genes per condition, roughly split between those whose loss caused depletion (sensitivity) or enrichment (resistance) (**Fig. 1I**). To avoid false positives from non-expressed genes, we filtered hits using A375 RNA-seq data from DepMap (Arafeh *et al*, 2025), which further refined and stratified the hit list (**Fig. 1J**). In summary, these screens provided a panoramic view of the genes that modulate the response to the NS-inducing agents ActD and BMH-21.

### A systems-level view of the pathways that modulate the response to NS

Beyond single genes, we performed a global analysis of the pathways that are involved in the response to ActD and BMH-21. First, and using the top and bottom 100 genes based on their expression levels for each treatment (TPM > 1; p < 0.01), we conducted a functional enrichment across Gene Ontology, Molecular Signatures Database, and Reactome pathways. We also annotated hits associated with the nucleolus (Human Cell Atlas (Thul *et al*, 2017)) or as cancer drivers (COSMIC Cancer Gene Census (Sondka *et al*, 2018)) (**Fig. 2A, Table S1**.).

**Figure 2.**
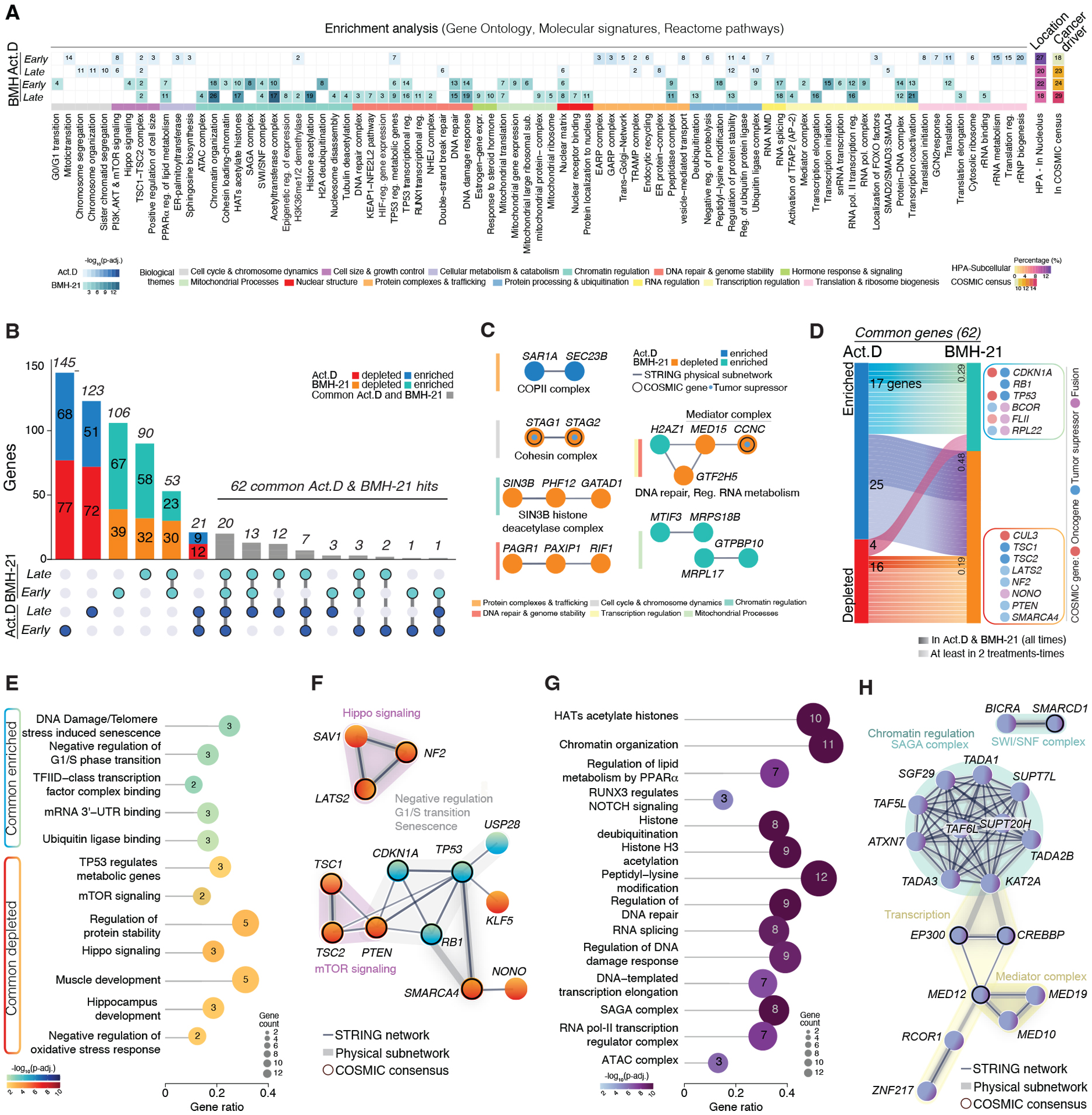
Comparative Functional and Network Analyses. (A) Functional enrichment analysis for the top and bottom 100 expressed genes (TPM > 1; p < 0.01) for each treatment (ActD, BMH-21) at early and late timepoints. To the right, the percentage of significant genes localized to the nucleolus (Human Protein Atlas Subcellular Compartment database) and the fraction classified in the COSMIC Cancer Gene Census (CGC) are shown, with absolute gene counts indicated within each cell. (B) UpSet plot illustrating the unique and shared top 100 significant genes across treatments and timepoints. Genes depleted upon ActD and BMH-21 treatment are shown in red and orange, while enriched genes are shown in blue and teal, respectively. Gray bars represent the 62 common top 100 significant genes shared between ActD and BMH-21 at different timepoints, highlighting their differential regulatory patterns. (C) STRING network analyses of interaction patterns among mutations unique to ActD or BMH-21. Thicker lines indicate higher confidence. Nodes with a black border denote COSMIC cancer driver genes, further annotated by their functional class (oncogene, tumor suppressor, or fusion gene). (D) Sankey plot depicting the 62 common top 100 significant genes from (B), indicating their roles as common sensitizers or resistance factors. Connections reflect whether genes display concordant (e.g., depleted in both treatments) or divergent regulatory patterns. Fractions indicate the proportion of common genes consistently observed across all conditions. COSMIC-listed genes are highlighted by their classification (oncogene, tumor suppressor, or fusion driver). (E) Enrichment analysis of pathways commonly enriched or depleted across ActD and BMH-21 treatments. Dot size reflects gene counts per term, and the −log10(adjusted p-value) indicates enrichment significance. Gene ratios (hits/total term genes) are annotated on the plots. (F) STRING network analyses of genes in (E) indicating key complexes including. Edges are scaled by STRING confidence, and grey sub-lines indicate experimentally validated physical interactions within a complex. Nodes with a black outline mark COSMIC cancer drivers, emphasizing their potential contribution to the enriched biological themes. (G) A lollipop enrichment plot illustrating the overrepresentation of chromatin regulators, transcriptional modulators, DNA repair factors, splicing components, and proteolytic machinery among genes enriched in ActD but depleted in BMH-21. Dot size corresponds to the number of genes, and −log10(adjusted p-value) reflects statistical significance. Gene ratios are annotated within the plot. (H) STRING network visualization of enriched chromatin remodeling complexes (SAGA, SWI/SNF), transcriptional regulators (Mediator complex), and DNA repair regulators among ActD-enriched/BMH-21-depleted genes. Edges are scaled by STRING confidence, with thicker lines denoting higher confidence of association. Grey sub-lines mark direct physical interactions within defined complexes. Nodes outlined in black denote COSMIC cancer driver genes, such as SMARCD1 (SWI/SNF), CREBBP, and MED12 (Mediator).

While some pathways were common across all treatments and timepoints (e.g. TSC1-TSC2, translation and ribosome biogenesis), these analyses evidenced that the response to these drugs differed significantly in many aspects. For instance, BMH-21 showed a specific enrichment for terms related to “protein–DNA complexes”, suggesting that part of its mechanism might involve direct protein-DNA adducts or regulating protein-DNA interactions. BMH-21 response genes were clearly enriched in DNA-damage-related and mitochondrial functions, including the mitochondrial ribosome and translation. Interestingly, ActD was enriched in pathways related to vesicular trafficking.

Next, we conducted a follow-up analysis using the top 100 significantly enriched and depleted genes across treatments. This comparison further highlighted that, while several mechanisms of response were shared between the drugs (62 genes in common), each compound also engaged distinct and non-overlapping pathways defining its specific response signature (**Fig. 2B**).

To gain mechanistic insight into these pathway signatures, we next mapped the significant hits onto STRING protein–protein interaction networks (Szklarczyk *et al*, 2023), focusing on validated physical interactions. This approach revealed coherent functional modules, where genes acting within the same molecular complexes or pathways clustered together. For instance, *STAG2* and *STAG1*, both established cancer drivers, emerged as a BMH-21-selective node, suggesting that cohesin integrity influences the response to this compound (**Fig. 2C**). This protein network analysis also highlighted the enrichment of epigenetic regulators and mitochondrial interactors among BMH-21 responsive genes, as well as the COPII vesicular traffic complex for ActD.

We then proceeded to interrogate the 62 common significant genes across both treatments. A Sankey plot illustrated the overlap between the two responses, distinguishing genes consistently enriched or depleted across both treatments from those driving drug-specific effects (**Fig. 2D**). Next, we focused on the mechanisms that are shared in the response to both drugs. Enrichment analyses and protein interaction analysis revealed that pathways related to G1/S transition, DNA damage-induced senescence, RNA polymerase II activity, RNA 3′ end regulation, and ubiquitin-mediated proteasomal degradation were related to resistance to both drugs, while those related to mTOR or Hippo signaling were related to sensitivity (**Fig. 2E, F**).

Finally, we examined genes that produced opposite responses to ActD and BMH-21. This analysis revealed a marked enrichment for chromatin regulators, transcriptional modulators, DNA repair factors, splicing components, and proteolytic machinery (**Fig. 2G, H**). Together, these analyses illustrate that while ActD and BMH-21 share key regulators of their sensitivity, they also differ significantly highlighting the need for ad hoc studies of drug-sensitivity for a better identification of biomarkers of response.

### Validation of hits that modify the sensitivity to ActD and BMH-21

To validate the screen findings, we performed cell-competition assays. A375 cells expressing Cas9 were partially infected with lentiviruses expressing an sgRNA under the U6 promoter and the Thy1.1 (CD90) ORF under EF1αs, enabling sgRNA tracking through CD90 surface expression (Umkehrer *et al*, 2021) (**Fig. 3A**). The frequency of CD90⁺ cells was monitored over time under each treatment, providing a readout of the impact of each mutation on drug response. As a control for this validation assay, we transduced cells with an sgRNA targeting the essential gene RPA3, observing the CD90⁺ population progressively depleted in the cultures irrespective of the treatment (**Fig. 3B).** Using this approach, we first confirmed that *TP53* deletion conferred resistance to both drugs (**Fig. 3C**). Unexpectedly, *STAG2* deletion increased sensitivity to ActD, not predicted from our screen, and particularly to BMH-21, for which it was predicted (**Fig. 3D**). Notably, these competition assays were performed using sgRNAs from a complementary CRISPR library, the Gecko library (Shalem *et al*, 2014). Nevertheless, this observation is of relevance since *STAG2* deletions are found in several cancers such as Ewing sarcoma, where ActD is part of the standard multiagent chemotherapy regimen (Jaffe *et al*, 1976), and in bladder cancer (Scott *et al*, 2025). Consistent with the screen data, targeting the SAGA complex factor TADA3 conferred resistance to ActD but sensitivity to BMH-21 (**Fig. 3E**), highlighting the mechanistic differences between both drugs. Testing independent sgRNAs for these factors, together with validation of on-target knockout by immunoblotting, confirmed the findings, including those for another SAGA core component, *SUPT20H* (**Fig. S2**). Overall, these analyses confirmed the results from our CRISPR screen and highlighted mutations that modulate the response to ActD and BMH-21.

**Figure 3.**
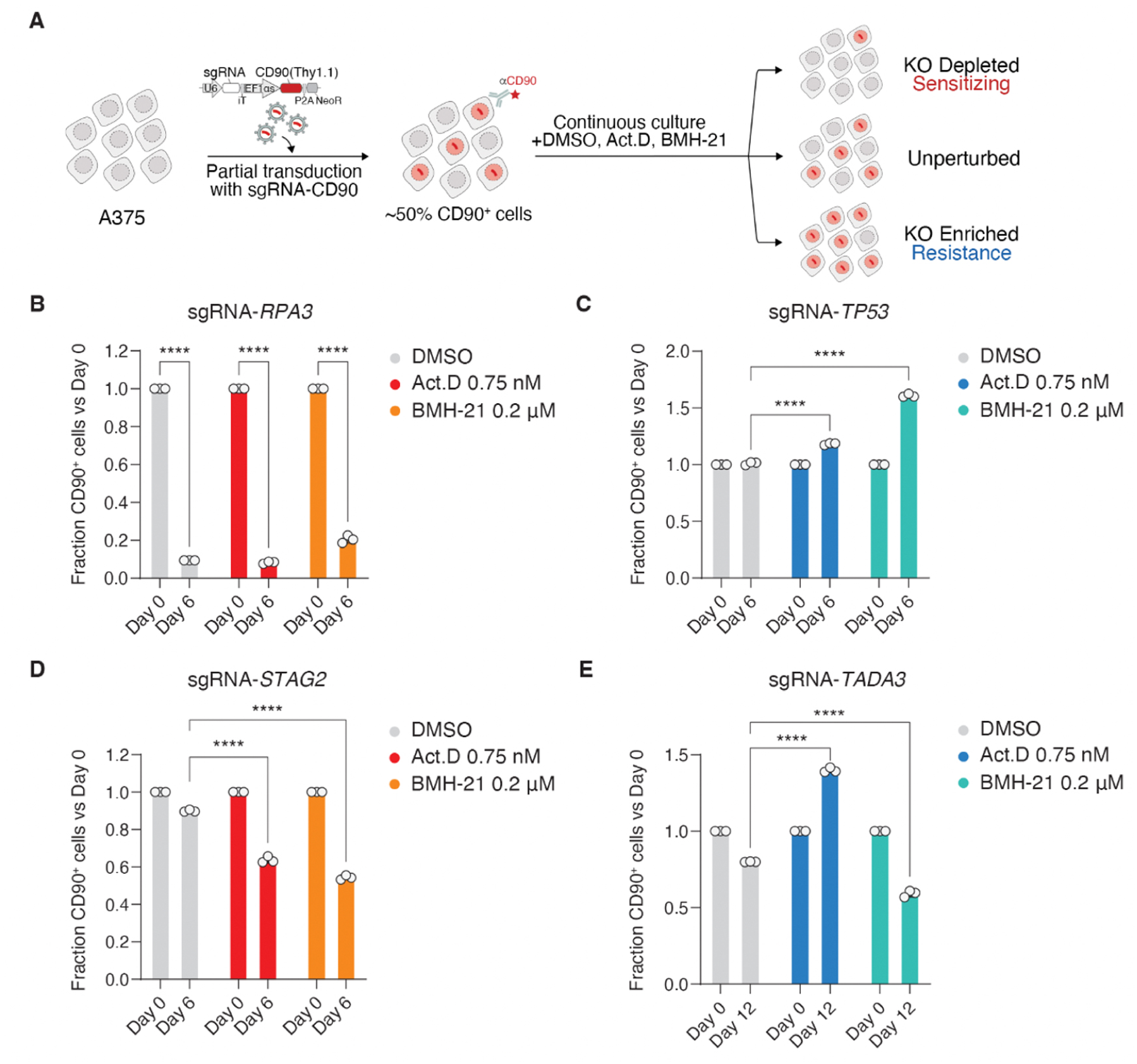
Hit validation by a competition assay. (A) Simplified overview of the competition assay. A375-Cas9-BFP cells from the CRISPR screen are partially infected by lentiviruses carrying a plasmid with the sgRNA sequence of the gene of interest and a CD90/Thy1.1 sequence, used as marker, leading to a pool of cells with ∼50% of cells carrying the plasmid. After continuous culture with ActD, BMH21 and DMSO the ratio of cells carrying the plasmid was evaluated over time. (B-E) Competition assay, as defined in A, using sgRNAs targeting *RPA3* (B), *TP53* (C), *STAG2* (D) and *TADA3* (E). ****, p < 0.0001

### PI3K/mTOR signaling shapes the response to therapies targeting RNA Pol I

Ultimately, our aim was to identify specific biomarkers that can predict the sensitivity to NS-inducing agents. From the different pathways identified in our screen, we focused on the mutations that enhanced PI3K and mTOR signaling, which are frequent events in cancer (Zhang *et al*, 2017). Unexpectedly, while PI3K and mTOR activity are typically associated with survival and proliferation, sgRNAs targeting *TSC1* and *TSC2* (which enhance mTOR signaling) or *PTEN* (which enhances PI3K signaling) increased the sensitivity to Act D and BMH-21 in the CD90 competition model defined above (**Fig. 4A** and **S3A, D**). Given the different dependencies shown by ActD and BMH-21, we wanted to explore the relationship between PI3K/mTOR and therapies targeting RNA Pol I by using a genetic system instead of drugs. To do so, we used a previously defined genetic model allowing conditional depletion of the Pol I catalytic subunit RPA194 (Ide *et al*, 2020). Here, doxycycline induces expression of the E3-ubiquitin ligase adaptor *TIR1*, and auxin promotes interaction between TIR1 and the degron-tagged RPA194, leading to its rapid proteasomal degradation (**Fig. 4B**). This system enabled a rapid and controlled depletion of RPA194, which translated into the loss of RNA Pol I activity, as seen by a reduction in the incorporation of 5’-Ethynyluridine (EU) (**Fig. 4C-E**). Importantly, deletion of *TSC1*, *TSC2* or *PTEN* enhanced the toxic effects of targeting RPA194 in this degron system (**Fig. 4E** and **Fig. S3E**).

**Figure 4.**
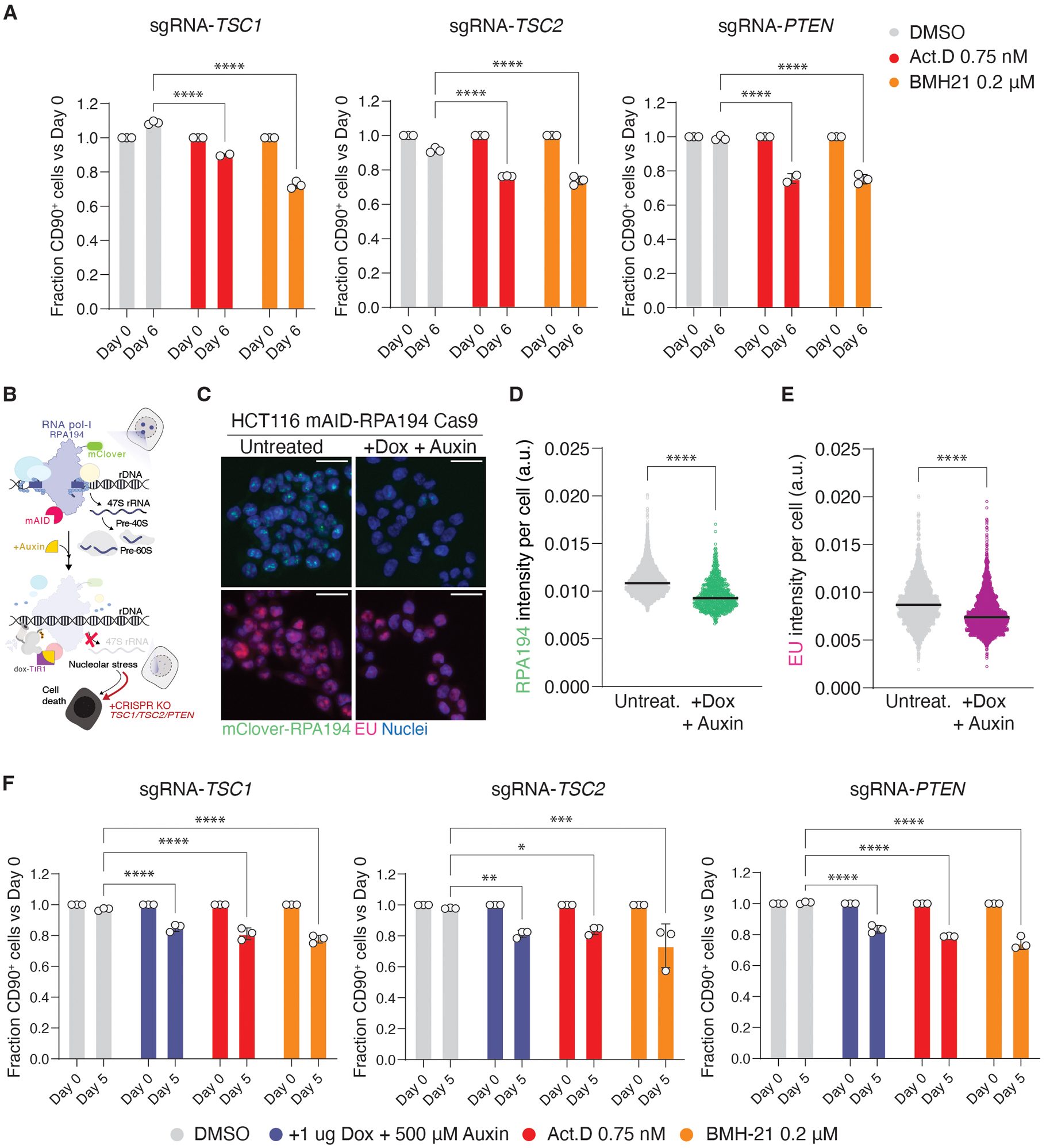
PTEN and TSC1/TSC2 modulate the response to therapies targeting RNA Pol. **I.** (A) Competition assay, as defined in Fig. 2A, using sgRNAs targeting *TSC1*, *TSC2* or *PTEN*. (B) Experimental design to assess whether loss of *TSC1*, *TSC2*, or *PTEN* enhances toxicity associated with nucleolar stress. The doxycycline- and auxin-inducible mAID-RPA194 system in HCT116 mAID-RPA194 Cas9 cells allows conditional degradation of RPA194: doxycycline induces TIR1 expression, and auxin promotes interaction between TIR1 and the degron-tagged RPA194, leading to its proteasomal degradation. (C) Immunofluorescence images illustrating the reduction in RPA194 and EU incorporation levels in HCT116 mAID-RPA194 Cas9 cells, upon exposure to doxycycline (1 μg) and auxin (500 μM) for 5h. Scale bar, indicates 30 μm. (D-E) High-Content Image analysis of the immunofluorescence data shown in (B), illustrating the reduction in RPA194 and EU levels in the in HCT116 cells expressing mAID-RPA194 upon treatment with doxycycline and auxin. Arbitrary units, a.u. (F) Competition assay performed in HCT116 cells expressing mAID-RPA194 using sgRNAs targeting *TSC1* (B), *TSC2* or *PTEN*. *, p < 0.05; **, p <0.01; *** p <0.001; ****, p < 0.0001

To further explore the impact of MTOR signaling in the NS response, we used *Tsc2*-deficient MEF. Of note, these MEF are also *p53*-deficient, as this condition is necessary for the growth of *Tsc2*-null cells. As observed in human cells, these MEF also presents sensitivity to BMH-21 and ActD (**Fig. 5A**). Furthermore, use of the mTOR inhibitor Rapamycin reduced the toxicity of ActD and BMH-21 in *Tsc2*-deficient MEF, further supporting that high levels of mTOR signaling sensitize to NS (**Fig. 5B, C**). Interestingly, Rapamycin alleviated the effect that these drugs have in the nucleolar area or in the redistribution of nucleolar factors such as FIBRILLARIN or NPM1, suggesting that the mechanism of our observation relates to a direct effect of PI3K/mTOR signaling in modulating nucleolar activity (**Fig. 5E, F**). Together, these experiments illustrate that mutations that enhance signaling through PI3K and mTOR, although they promote the survival and growth of cancer cells, they render them sensitive to therapies that target RNA Pol I.

**Figure 5.**
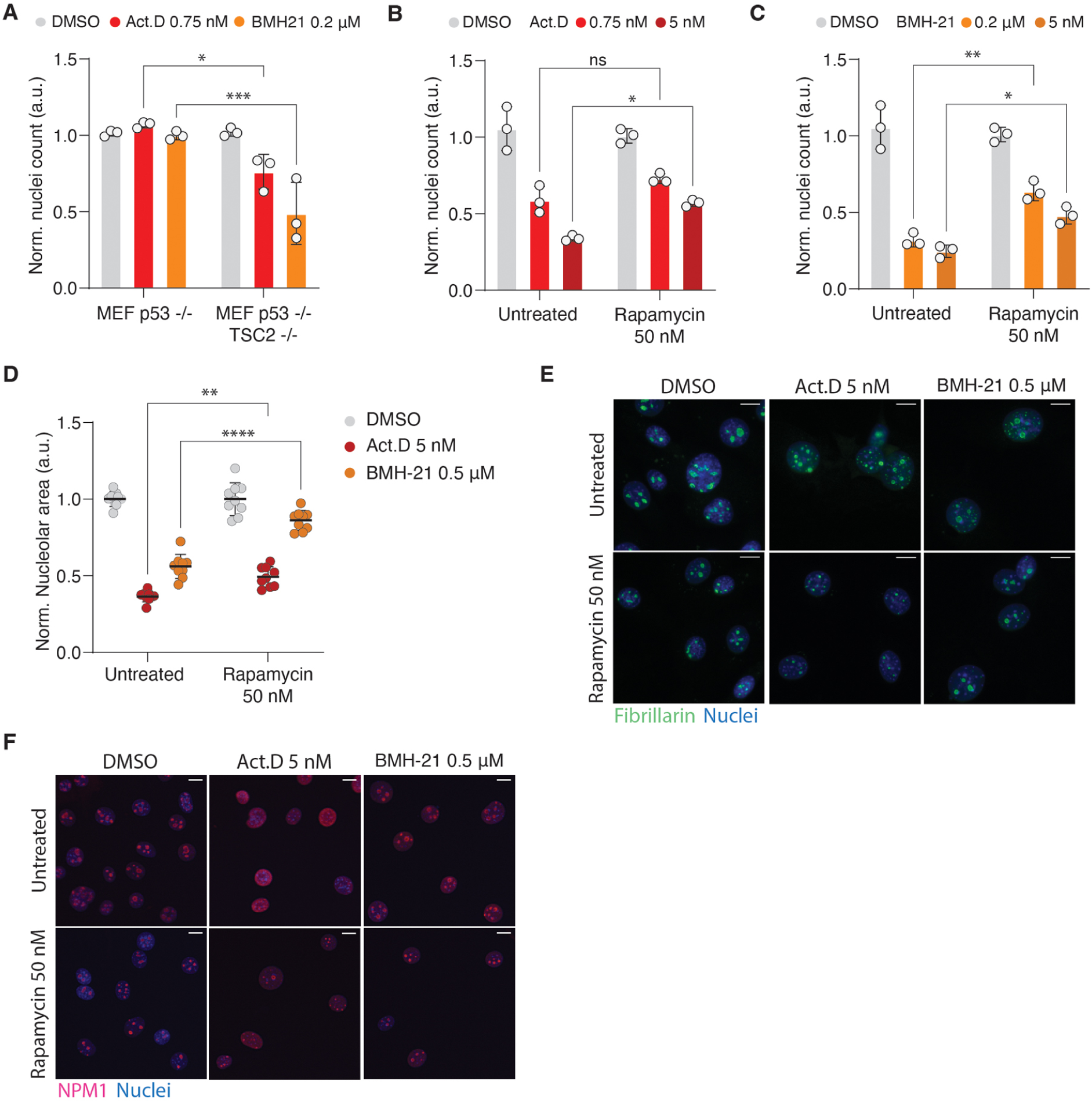
mTOR inhibition protects cells from NS. (A) Nuclei counts, as quantified by High Content Microscopy, in *p53*^−/−^*Tsc2*^+/+^ and *p53*^−/−^*Tsc2*^−/−^ MEF grown in the presence or absence of the indicated drugs. Nuclei counts were normalized to those found on the control condition (DMSO). (B, C) Nuclei counts, as quantified by High Content Microscopy, in NIH3T3 cells exposed to BMH-21 or ActD, upon a previous exposure to rapamycin for 24h. (D) High Content Microscopy-mediated quantification of nucleolar area (based on FIBRILLARIN staining) in NIH3T3 cells treated as mentioned in panels B, C. (E) Representative images of the FIBRILLARIN staining in NIH3T3 from the analysis shown in Fig. 5D. Scale bar, indicates 2 μm. (F) Representative images of redistribution of NPM1 in NIH3T3 cells treated as in Fig. 5 (B, C). Scale bar, indicates 2 μm. ns, not significant; *, p < 0.05; **, p <0.01; *** p <0.001; ****, p < 0.0001. Arbitrary units, a.u.

## DISCUSSION

Our original intention was to better understand the mechanisms that shape the response to nucleoli-targeting drugs, as these drugs are now entering clinical development, and a better stratification of the patients would be necessary to design the clinical trials. The work presented here has identified several genetic biomarkers that predict the sensitivity or resistance to these agents. Among these, mutations in key tumor suppressors such as *TP53*, as well as components of the SAGA complex and genes associated with the PI3K/mTOR signaling pathway (e.g., *PTEN, TSC1, TSC2*), emerge as important modulators of the response to NS. Importantly, mutations that enhance PI3K/mTOR signaling, which are frequently observed across diverse cancer types, render cells more susceptible to genetic or chemical therapies targeting RNA Pol I, establishing these mutations as potential predictive biomarkers for therapeutic response. Significantly, the activity of PI3K/mTOR signaling can be easily monitored in human cancer biopsies, supporting the feasibility of this biomarker.

A striking observation arising from our study is the differential—and sometimes even opposing—response conferred by certain mutations to ActD and BMH-21. Although both drugs are thought to be toxic due to their effects in inhibiting RNA Pol I activity, our data demonstrate that some genetic lesions, such as those affecting the SAGA complex, confer resistance to one while sensitizing cells to the other. These data raise a word of caution in the development of therapies targeting the nucleoli or RNA Pol I, since an in-depth understanding of their mechanism would be necessary to guide their clinical use in a reasoned manner.

From a translational perspective, our findings might have clinical implications. By defining a global landscape of genetic vulnerabilities and resistance mechanisms to NS, our work provides a rationale for using these biomarkers to guide patient stratification in clinical trials of RNA Pol I inhibitors. Identifying tumors with enhanced PI3K/mTOR signaling or altered SAGA complex function may help prioritize patients who stand to benefit most from this therapeutic strategy, while sparing others from ineffective and potentially toxic treatments.

Finally, our study underscores the importance of performing an in-depth molecular phenotyping for oncology drugs that are progressing to clinical development. There is a rising awareness that drugs have multiple targets and that their off-targets might significantly contribute to their anticancer effects (Lin *et al*, 2019). Our work further underscores the need of a mechanism-aware approach to cancer therapy.

## MATERIALS AND METHODS

### Cell lines

Melanoma A375 Cas9-BFP cells were kindly provided by CRISPR Functional Genomics (CFG) facility at Karolinska Institutet and cultured in DMEM supplemented with 10% fetal bovine serum (FBS) and 1 % penicillin-streptomycin (PE/ST). HEK293T cells (ATCC CRL-11268) were used for lentiviral production and cultured under the same conditions. HCT116 mAID RPA194 mClover cells were provided by Kazuhiro Maeshima and Satoru Ide (Ide *et al*., 2020) and maintained in McCoy’s 5A medium supplemented with 10% FBS and 1% PE/ST. To generate HCT116 mAID-RPA194-Cas9 cells, the parental line was transduced with a lentivirus encoding BFP-tagged Cas9 (Addgene #196714, provided by CFG). Transduced cells were selected with 10 µg/mL blasticidin (Thermo Fisher, R21001) for 10 days, followed by clonal expansion. Stable Cas9 expression was confirmed by immunofluorescence (IF) and western blot (WB). NIH3T3 cells were obtained from CNIO. p53-deficient and Tsc2-deficient mouse embryonic fibroblasts (MEFs) were kindly provided by Manuel Serrano and maintained in DMEM supplemented with 15% FBS and 1% PE/ST.

### Genome-wide pooled CRISPR screen

The A375 melanoma cell line was engineered to stably express S. pyogenes Cas9. A lentiviral vector encoding Cas9, blasticidin resistance, and BFP (Addgene #196714) was transduced, and after blasticidin selection, BFP-high cells were twice sorted by FACS, expanded, and used for the genome-wide CRISPR screen.

The Brunello sgRNA library (Doench *et al*., 2016), targeting 19,114 genes with four sgRNAs per gene and 1,000 non-targeting controls, was resynthesized with Unique Molecular Identifiers (UMIs) (Schmierer *et al*, 2017) to allow precise PCR-based quantification and amplification. sgRNAs were synthesized as pooled oligos (CustomArray), cloned into a modified lentiGuide-Puro backbone (Addgene #52963) carrying an AU-flip element (Cross *et al*, 2016), and packaged into lentivirus.

A375 Cas9-BFP cells were transduced in duplicate at a low multiplicity of infection (MOI ≈ 0.4) to ensure predominantly single sgRNA integration, maintaining ∼1,000× library coverage. Transductions were performed in the presence of polybrene (2 µg/mL), and after 48 h, puromycin (2 µg/mL; Jena Bioscience NU-931-05) was applied for two days to remove untransduced cells. After selection, 80 million cells per replicate were harvested as the T0 reference population, preserving full library representation. The remaining cells were seeded into T175 flasks (∼6 million cells per flask), maintaining 80 million cells per replicate for each treatment condition.

Cells were allowed to attach for 24 h before replacing the medium with DMEM containing either DMSO (control), Actinomycin D (ActD, 0.75 nM; Sigma A1410-10MG), or BMH-21 (0.2 µM; Sigma SML1183-5MG). After three days of treatment, cells were counted, and 80 million were collected per replicate as the first treatment timepoint. In parallel, another 80 million cells per condition were reseeded in fresh compound-containing media, maintaining continuous exposure. This collection–reseeding cycle was repeated every three days until day 12.

Early timepoints corresponded to approximately 50% growth inhibition (IC₅₀), reached on day 3 for ActD and day 6 for BMH-21, capturing genes whose loss rapidly sensitized cells to nucleolar stress. The late timepoint, day 12, was used to identify genes conferring longer-term resistance or adaptation.

Genomic DNA was extracted from all collected pellets and from the original plasmid library using the QIAamp DNA Blood Maxi Kit (Qiagen #51192). Guide and UMI sequences were PCR-amplified and prepared for next-generation sequencing following (Schmierer *et al*., 2017).

### Computational analyses of genome-scale CRISPR screens

Next-generation sequencing reads were processed using the MaGeCK software (Li *et al*., 2014) combined with UMI lineage dropout analysis (Schmierer *et al*., 2017). sgRNA counts for each treatment and timepoint were normalized to the corresponding DMSO controls. Replicates were averaged, producing a single log2 fold-change (log2FC) value for each sgRNA, and gene-level scores were derived by averaging all sgRNAs targeting the same gene. Gene-level significance and ranking were determined using MaGeCK robust rank aggregation (RRA) to generate p-values for both depletion and enrichment. To enable direct comparison across directions of effect, we applied a unified ranking scheme: negative log₂FC values retained negative RRA scores (sensitizers), while positive log₂FC values retained positive scores (resistance genes). This produced a single continuous ranking, where lower ranks indicated stronger depletion and higher ranks stronger enrichment. Genes with an RRA p-value < 0.01 were considered significant hits. To remove potentially non-relevant library artifacts, the expression of these genes was validated using batch-corrected RNA-seq expression data for A375 cells from the Broad DepMap dataset (batch-corrected DepMap 24Q4) (Tsherniak *et al*, 2017), excluding genes with TPM < 1. These results at gene and sgRNA level are summarized on **Table S1**. All data visualization was performed in RStudio using ggplot2 v3.5.1 (Wickham, https://cran.r-project.org/web/packages/ggplot2/index.html), UpSetR v1.4.0 ((Conway *et al*, 2017), https://cran.rstudio.com/web/packages/UpSetR/), and pheatmap (Kolde, https://github.com/raivokolde/pheatmap).

### Enrichment and network analysis of CRISPR screen hits

For each treatment condition, gene set enrichment analysis was performed on the top 100 significantly enriched and depleted hits (RRA p < 0.01, TPM > 1). Gene symbols were mapped to Entrez IDs using Brunello annotations, and enrichment was assessed against Gene Ontology (biological process, molecular function, and cellular component) (Gene Ontology *et al*, 2023), Reactome pathways (Milacic *et al*, 2024), and Hallmark gene sets from MSigDB (Liberzon *et al*, 2015) using ClusterProfiler (Wu *et al*, 2021). Benjamini–Hochberg correction was applied to obtain adjusted p-values, retaining only terms with at least two genes. To improve interpretability, the top 10 most significant and nonredundant terms were selected for visualization per condition and direction. Results from enrichment analysis are summarized on **Table S1**.

Protein–protein interaction networks were generated with STRING (Szklarczyk *et al*., 2023), including both complete interaction networks and high-confidence physical subnetworks (STRING score > 0.7). In these networks, edge thickness corresponds to STRING confidence scores, and additional grey sub-lines indicate experimentally validated physical interactions within complexes.

Finally, significant hits were annotated for protein localization to the nucleolus using the Human Protein Atlas subcellular compartment database (Thul *et al*., 2017) and cross-referenced with the COSMIC Cancer Gene Census (Sondka *et al*., 2018), which also provided functional classifications of oncogenes, tumor suppressors, and fusion drivers.

### Guide cloning for competition assays

Guide sequences used for validation were obtained from the GeCKO library (Shalem *et al*., 2014) and are listed in **Table S2**. Guide cloning was performed according to standard protocols. Briefly, top- and bottom-strand oligos (Eurofins) were resuspended in water at 100 mM and then mixed at 1:1 ratio for each sgRNA. Then, 1 µL of the oligo mix was added to a master mix containing 1× T4 ligase buffer (NEB M0202M) and 0.5 µL T4 PNK (New England Biolabs (NEB), M0201S), and water to a final volume of 10 µL per reaction. For oligo annealing, we incubated the oligo mix at 37°C for 30 min, then 95°C for 5 min with a temperature change of 1°C every 5 s until reaching 25 °C. Then, the annealed oligos were diluted 1:100 and cloned into hU6-sgRNA-EFIas-Thy1.1-P2A-NeoR, kindly obtained by Johannes Zuber (Umkehrer *et al*., 2021), using the NEBridge Golden Gate Assembly (BsmBI-v2) Kit (NEB, E1601). Plasmids were expanded on *E. coli* Stellar competent cells (Takara) grown on 100 µg/mL Carbenicillin (Fisher Scientific, 11480952). Guide plasmids were isolated using QIAprep Spin Miniprep Kit (Qiagen, 27104), and guide inserts were verified by Sanger sequencing (Microsynth).

### Lentiviral production

Lentiviral supernatant was produced in HEK293T cells (ATCC CRL-11268) by reverse transfection with Lipofectamine 3000 (Fisher Scientific, L3000008), pMD2.G (Addgene, no.12259), psPAX2 (Addgene no.12260) plasmids, and the transfer plasmids expressing the sgRNA sequence and the CD90/Thy1.1 reporter. After 24 h, the medium was replaced, and virus-rich supernatants were collected 48 h post-transfection, filtered, and stored at −80 °C.

### Competition Assay

A375 Cas9-BFP cells and HCT116 mAID-RPA194-Cas9 cells were seeded in 6-well culture plates (100 000 cells/well) and infected with lentiviral supernatants containing sgRNA–Thy1.1 constructs to achieve 30–60% transduction efficiency. The following day, cells were washed several times with PBS and reseeded into fresh 6-well plates. After an additional 24 h, cells were collected and replated at 80 000 cells/well for the start of the assay.

For A375-Cas9-BFP cells, treatments began immediately (day 0) in medium containing Actinomycin D (0.75 nM), BMH-21 (0.2 µM), or DMSO. HCT116 mAID-RPA194-Cas9 cells were pre-treated with 1 µg/mL doxycycline for 24 h to induce TIR1 expression. On the next day (day 0), cells were treated with ActD (0.75 nM), BMH-21 (0.2 µM), DMSO, or a combination of doxycycline (1 µg/mL) and auxin (500 µM) to induce RPA194 degradation.

A375 cells were split and re-treated every three days, while HCT116 cells were split every two days. At day 0 and each split point, cells were collected and analyzed by flow cytometry (Guava easyCyte HT) to quantify the proportion of CD90⁺ cells, representing the sgRNA-expressing population. Cells were stained with anti-CD90/Thy1.1-FITC (Thermo Fisher 11-0900-81, 1:4000 dilution in 2% FBS/PBS) for 1 h at 4 °C in the dark, washed twice with PBS, and resuspended in 2% FBS/PBS for analysis.

To verify knockout efficiency, sgRNA-expressing A375 and HCT116 cells were selected with 900 µg/mL Geneticin (Thermo Fisher 10131027) for 7 days, collected, and analyzed by western blot.

### Immunofluorescence and image analysis

Throughout this work, we followed this standard protocol. Cells seeded and cultured onto 96-well imaging plates. On the experiment endpoint, cells were fixed with 4% PFA for 15 min and permeabilized with 0.1% Triton X-100 (Sigma Aldrich, X100) for 10 min at room temperature (RT). After blocking (3% BSA and 0.1% Tween-20 in PBS) for 30 min at RT, cells were incubated with primary antibody in the same blocking solution overnight at 4°C. The indicated concentration of primary antibodies was applied: anti-Cas9 (Abcam, ab191468, 1:1000), anti-Fibrillarin (Abcam, ab5821, 1:1000), anti-NPM1 (Abcam, ab10530, 1:500), and anti-GFP primary antibody (Abcam, ab290, 1:2000) was used to boost the signal from mClover tagged RPA194 in HCT116 mAID RPA194 Cas9 cells, as mClover is a derivative from GFP. The next day, cells were washed again with PBS and incubated with secondary antibody in blocking solution at a 1:500 dilution (either Alexa Fluor 488 conjugated secondary antibody (Thermo Fisher, A-11008) or Alexa Fluor 647 conjugated antibody (Thermo Fisher, A-21235)), and were kept in the dark for 1 h at RT. Next, the cells were washed and their nuclei were stained with 2 μM Hoechst 33342 (Sigma Aldrich, 14433) for 10 min. Then cells were washed in PBS and imaged. Images were acquired using an IN Cell Analyzer 2200 (GE Healthcare, USA), and quantitative image analyses were run in the open-source software CellProfiler (www.cellprofiler.org) (Jones *et al*, 2008). We used self-made image analysis pipeline segmenting nuclei based on Hoechst signal and then measuring the intensity of the marker of interest within the nucleus, such as for RPA194 or EU measurements. Within the nucleus, the pipeline can segment nucleoli, based on expression of nucleolar markers such as Fibrillarin, enabling automated measurement of changes in nucleoli area.

### EU RNA labeling

HCT116 mAID-RPA194-Cas9 cells were seeded in 96-well imaging plates (BD Falcon 353219) at 4,000 cells per well. The following day, cells were treated with 1 µg/mL doxycycline (Sigma Aldrich C8895) for 24 h to induce TIR1 expression. The medium was then replaced with fresh medium containing 1 µg/mL doxycycline and 500 µM auxin (Sigma Aldrich I3750-5G-A), while control wells received medium only. After 4 h, 5-ethynyl uridine (EU) was added to a final concentration of 1 µM (Click-iT RNA Imaging Kit, Invitrogen C10329) and cells were incubated for 1 h at 37 °C to allow incorporation into nascent RNA, primarily rRNA. Cells were then fixed with 4% formaldehyde, and EU detection was performed according to the manufacturer’s protocol. Nuclei were counterstained with Hoechst, and plates were imaged and analyzed as described above.

### Cell viability assays

To assess the sensitivity of Tsc2-deficient cells to RNA Pol I inhibition, p53⁻/⁻ and p53⁻/⁻ Tsc2⁻/⁻ MEFs were seeded in 96-well imaging plates (BD Falcon 353219) at 1,000 cells per well. The following day, cells were treated with Actinomycin D (0.75 nM or 5 nM), BMH-21 (0.2 µM or 0.5 µM), or DMSO as control. After 48 h, plates were fixed with 4% formaldehyde (Sigma Aldrich F8775-500ML), nuclei were stained with Hoechst, and images were acquired and analyzed as described above.

### Assessment of mTOR inhibition on Pol I stress response

To test whether mTOR inhibition protects against Pol I–targeting compounds, NIH3T3 cells were seeded in 96-well imaging plates (BD Falcon 353219). The following day, cells were treated with 50 nM rapamycin (Sigma Aldrich 37094) or left untreated. After 24 h, the medium was replaced with fresh medium containing either no treatment or rapamycin, and cells were co-treated with Actinomycin D (0.75 nM or 5 nM), BMH-21 (0.2 µM or 0.5 µM), or DMSO. After 48 h, plates were processed for immunofluorescence staining as described above.

### Western blotting

Cells were washed with PBS upon trypsination (Thermo Fisher, 15400054) and resuspended in RIPA buffer (Thermo Fisher, PI-89901) on ice. After sonication and centrifugation at 16,900 x g for 20 min at 4°C the supernatant was collected, and the protein concentration was determined using the DC Protein Assay Kit II (Bio-Rad, 5000112). The samples were heated at 95 °C for 5 min in a mixture of NuPAGE LDS Sample Buffer (Invitrogen, NP0007) and NuPAGE Reducing Agent (Invitrogen, NP0009). Electrophoresis was carried out on Bis-Tris gels and blotted on nitrocellulose membranes from Bio-Rad (1704270). Membranes were blocked in 5% milk, diluted in TBS-Tween buffer, followed by an overnight incubation at 4 °C with primary antibody: anti-Cas9 antibody (Abcam, ab191468, 1:1000), p53 (Abcam, ab1101, 1:1000), STAG2 (D25A4) (Cell Signaling Technology, 5882, 1:1000), TADA3 (Atlas Antibodies, HPA042250, 1:500), Hamartin/TSC1 (D43E2) (Cell Signaling Technology, 6935, 1:500), Tuberin/TSC2 (D93F12) XP (Cell Signaling Technology, 4308, 1:500), PTEN (D3Q6G) (Cell Signaling Technology, 14642, 1:1000), GAPDH (Abcam, ab9485, 1:2500) (also summarized on **Table S2**). After primary incubation, the membranes were washed with TBS-Tween buffer and incubated with secondary antibody for 1 h at RT. Protein bands were visualized by chemiluminescence (either with ECL, Thermo Scientific, 34076, or Amersham ECL, GE Healthcare, RPN2235) and imaged on an Amersham Imager 600 (GE Healthcare, USA).

### Statistics

Statistical analyses were carried out using GraphPad Prism (GraphPad Software Inc, version 10.5.0). To compare multiple data points with each other within an experimental condition, two-way-ANOVA tests were performed. In experiments where only two data points were compared, the Mann Whitney test was used.

## Supporting information

Table S1: CRISPR Screen; raw data

Table S1: List of reagents

## ACKNOWLEDGEMENTS

We thank the CRISPR Functional genomics (CFG) facility at Karolinska Institutet, and specially Jenna Persson and Bernhard Schmierer for their help with the CRISPR screen performance and analysis. Research was funded by grants from the Cancerfonden foundation (CAN 24/3438) and the Swedish Research Council (VR) (538-2014-31) to O.F-C.

## AUTHOR CONTRIBUTIONS

M.B. contributed to experiments related to validation and cellular analyses, preparation of the figures and to writing the manuscript. A.C. conducted the CRISPR screen, its bioinformatic analysis, and preparation of the figures. M.H. and L.L. provided experimental support to cell and molecular biology experiments. A.G. and M.M. helped with analyses related to TSC complex and mTOR. D.H. helped to coordinate the study and supervise the work. A.C. and O.F-C. supervised the study and wrote the manuscript.

## DECLARATION OF INTERESTS

The authors declare no competing interests.

## Notes

### Competing Interest Statement

The authors have declared no competing interest.

