## Supplemental Figs 1-3 for "PI3K/mTOR activity sensitizes cancer cells to nucleolar stress"

**Figure S1. Quality control of the genome-wide nucleolar stress CRISPR screen. *Related to Figure 1*.**

(A) Brunello-UMI lentiviral plasmid design. Schematic of the lentiviral backbone used to deliver the genome-wide Brunello library, targeting 19,114 genes with 4 sgRNAs per target plus 1,000 non-targeting (NT) controls. Each sgRNA is flanked by an Illumina i7 adapter sequence and a unique molecular identifier (UMI) to enable accurate PCR amplification and quantitative deconvolution of sgRNA abundances after next-generation sequencing.

(B) Distribution and representation of sgRNAs in the plasmid library. Next-generation sequencing of the plasmid pool confirmed uniform representation of sgRNAs, with a 90:10 skew ratio of 2.63, indicating no major bias. Importantly, 99.98% of all sgRNAs were detected, and 100% of gene targets were represented, confirming the integrity and evenness of the starting library prior to lentiviral packaging and transduction.

(C) Reproducibility of sgRNA abundance across replicates. Pearson's correlation heatmaps show pairwise correlations between biological replicates (R1, R2) for all conditions (untreated, DMSO, Act.D, and BMH-21) at early and late timepoints. High correlation coefficients ($r>0.9$ for most conditions) indicate robust reproducibility of sgRNA representation across replicates and treatment conditions.

(D) Baseline behavior of non-targeting (NT) and gene-targeting (GT) sgRNAs. Enrichment plots show log2 fold-change (log2FC) distributions of NT vs GT sgRNAs for all conditions relative to the untreated T0 reference population. NT guides remain centered around log2FC≈0, whereas GT guides display broader distributions under selective pressure, confirming that depletion and enrichment effects reflect true biological selection rather than technical noise.

(E) Validation of screen hits outside the main figure. Strip plots show log2FC values for all sgRNAs targeting the top 10 enriched and depleted genes from Figure 1D–G that were not included in Figure 1H. Filled colored squares show the mean log2FC per target gene under each treatment, while grey squares represent the DMSO baseline. The 95% confidence interval of NT sgRNAs is shaded to highlight the neutral control distribution. These data reinforce the consistency of sgRNA-level effects for the strongest hits across treatments and timepoints.

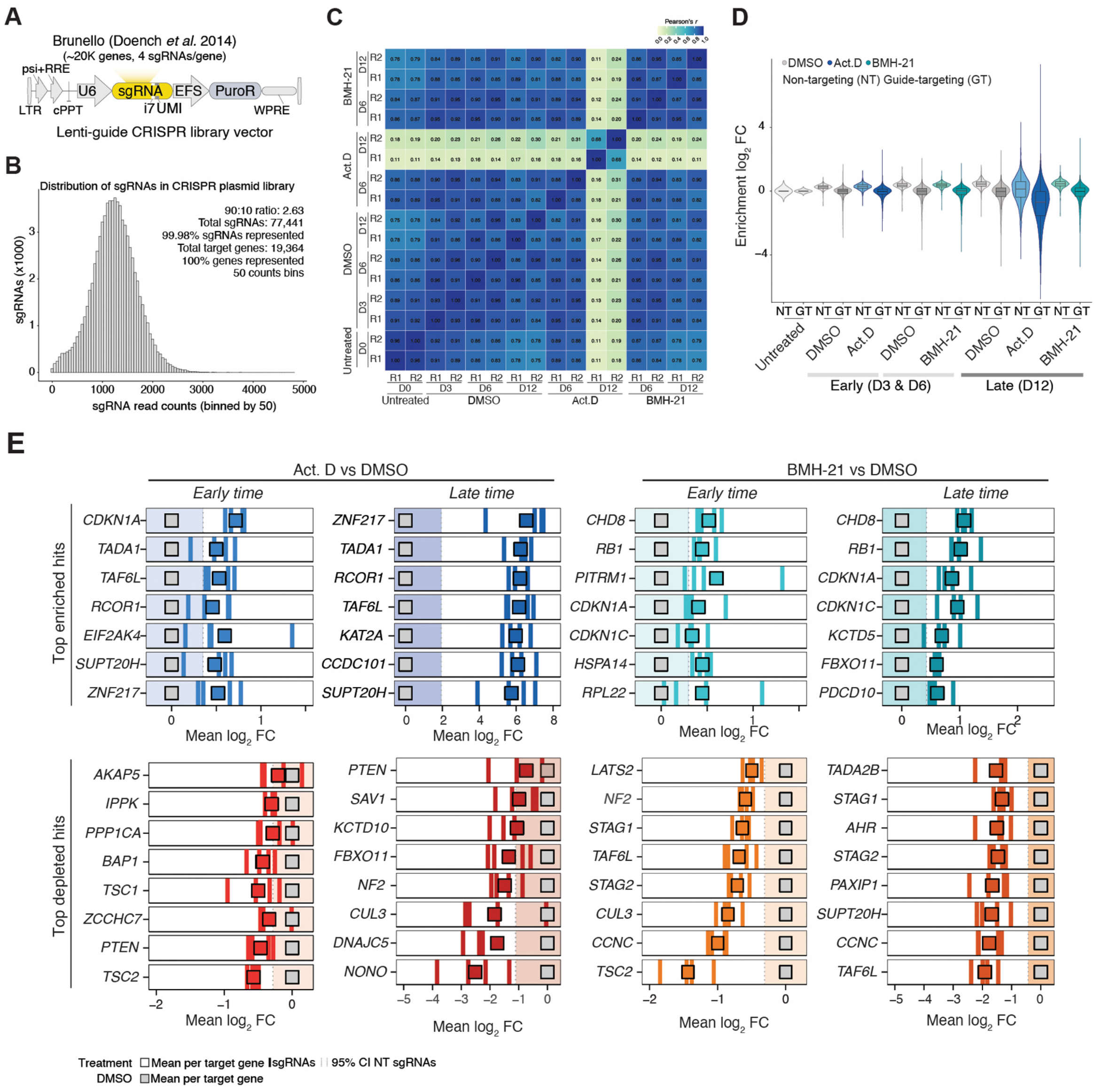
A
Brunello (Doench et al. 2014)
(~20K genes, 4 sgRNAs/gene)
psi+RRE
U6
sgRNA
EFS
PuroR
LTR
cPPT
i7UMI
WPRE
Lenti-guide CRISPR library vector
B
Distribution of sgRNAs in CRISPR plasmid library
90:10 ratio: 2.63
Total sgRNAs: 77,441
99.98% sgRNAs represented
Total target genes: 19,364
100% genes represented
50 counts bins
sgRNAs (x1000)
sgRNA read counts (binned by 50)
C
Pearson's r
BMH-21
Act.D
DMSO
Untreated
D0
D3
D6
D12
R1
R2
D
DMSO
Act.D
BMH-21
Non-targeting (NT) Guide-targeting (GT)
Enrichment log2 FC
NT GT
Untreated
Early (D3 & D6)
Late (D12)
E
Act. D vs DMSO
BMH-21 vs DMSO
Early time
Late time
Top enriched hits
Top depleted hits
CDKN1A
TADA1
TAF6L
RCOR1
EIF2AK4
SUPT20H
ZNF217
KAT2A
CCDC101
CHD8
RB1
PITRM1
CDKN1C
HSPA14
RPL22
KCTD5
FBXO11
PDCD10
AKAP5
IPPK
PPP1CA
BAP1
TSC1
ZCCHC7
PTEN
TSC2
SAV1
KCTD10
NF2
CUL3
DNAJC5
NONO
LATS2
STAG1
STAG2
CCNC
TADA2B
AHR
PAXIP1
Mean log2 FC
Treatment
DMSO
Mean per target gene
sgRNAs
95% CI NT sgRNAs

**Fig. S2. Additional validation experiments for hits from the CRISPR screen. *Related to Figure 3*.**

(A-C) Western blot showing the reduction of the proteins p53, STAG2 and TADA3 in A375-Cas9-BFP cells carrying sgRNA sequences for TP53, STAG2 and TADA3. For each gene two sgRNA sequences were tested (#1 and #2). GAPDH levels are shown as a loading control.

(D-H) Competition assay, as defined in Fig. 2A, using additional sgRNAs targeting *RPA3, TP53, STAG2, TADA3* and *SUPT20H*.

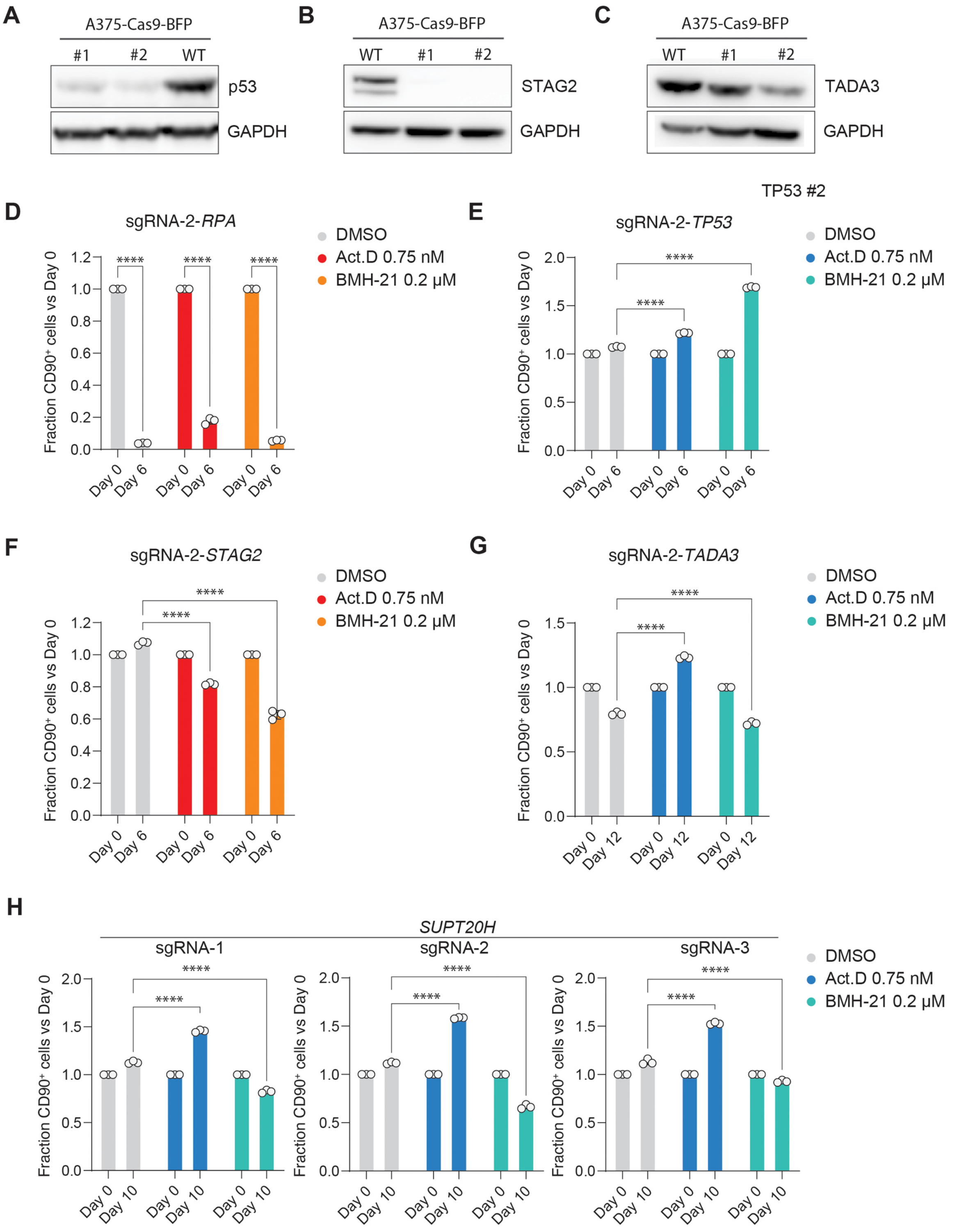
A
A375-Cas9-BFP
#1 #2 WT
p53
GAPDH
B
A375-Cas9-BFP
WT #1 #2
STAG2
GAPDH
C
A375-Cas9-BFP
WT #1 #2
TADA3
GAPDH
TP53 #2
D
sgRNA-2-RPA
DMSO
Act.D 0.75 nM
BMH-21 0.2 µM
Fraction CD90+ cells vs Day 0
Day 0
Day 6
****
E
sgRNA-2-TP53
DMSO
Act.D 0.75 nM
BMH-21 0.2 µM
F
sgRNA-2-STAG2
G
sgRNA-2-TADA3
Day 12
H
SUPT20H
sgRNA-1
sgRNA-2
sgRNA-3
Day 10

**Fig. S3. Additional analyses related to PTEN and the TSC complex.** ***Related to Figure 4*.**

(A) Western blot showing the reduction of the proteins TSC1, TSC2 and PTEN in A375-Cas9-BFP cells carrying the indicated sgRNA sequences. GAPDH levels are shown as a loading control.

(B-D) Competition assay, as defined in Fig. 2A, using additional sgRNAs targeting *TSC1, TSC2* and *PTEN*.

(A) Western blot showing the reduction of the proteins TSC1, TSC2 and PTEN in HCT116 cells expressing mAID-RPA194, Cas9, and sgRNAs targeting these genes. GAPDH levels are shown as a loading control.

Barz *et al* **Fig S3**

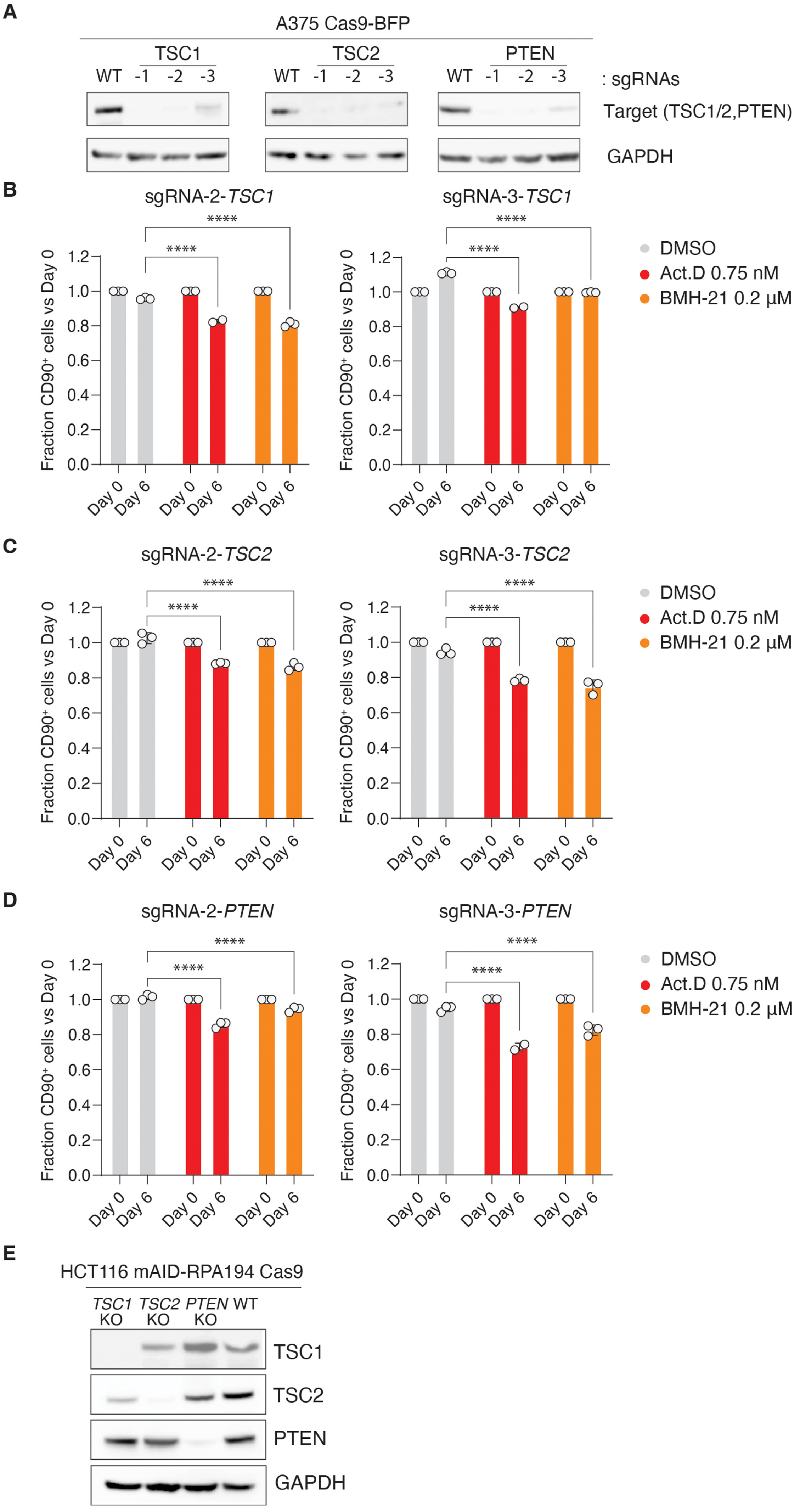
