## Supplementary material for "PI3K/mTOR activity sensitizes cancer cells to nucleolar stress": Table S1: List of reagents

**Table S2. List of reagents used in this study.**

| Cell lines | Species | Source |
|---|---|---|
| A375-Cas9-BFP | Human | CRISPR Functional Genomics facility (KI) |
| HCT116 mAID RPA194 mClover | Human | Satoru Ide et al, 2020 |
| MEF p53 -/- | Mouse | Manuel Serrano |
| MEF p53 -/- TSC2 -/- | Mouse | Manuel Serrano |
| NIH3T3 | Mouse | From CNIO |
| HEK293T | Human | ATCC, CRL-11268 |

| Plasmid | Source | Selection marker |
|---|---|---|
| p23_Cas9-BFP | CRISPR Functional Genomics (CFG) facility | Blasticidin |
| pMD2.G | Addgene (no. 12259) | - |
| psPAX2 | Addgene (no. 12260) | - |
| hU6-sgRNA-EFlas-Thy1.1-P2A-NeoR | Kind gift from Johannes Zuber. | Neomycin (Geneticin) |

| Antibodies | Source | Catalogue No | Application |
|---|---|---|---|
| Cas9 | Abcam | ab191468 | WB/IF (1:1000) |
| p53 | Abcam | ab1101 | WB (1:1000) |
| STAG2 (D25A4) | Cell Signaling technology | 5882 | WB (1:1000) |
| TADA3 | Atlas Antibodies | HPA042250 | WB (1:500) |
| Hamartin/TSC1 (D43E2) | Cell Signaling technology | 6935 | WB (1:500) |
| Tuberin/TSC2 (D93F12) XP | Cell Signaling technology | 4308 | WB (1:500) |
| PTEN (D3Q6G) | Cell Signaling technology | 14642 | WB (1:1000) |
| GAPDH | Abcam | ab9485 | WB (1:2500) |
| Cd90 (Thy 1.1) Monoclonal Antibody (HIS51), FITC | Thermo Fisher | 11-0900-81 | FACS (1:4166) |
| Cd90 Monoclonal Antibody (5E10), PE | Thermo Fisher | A15794 | FACS (1:4166) |
| GFP | Abcam | ab290 | IF (1:2000) |
| Fibrillarin | Abcam | ab5821 | IF (1:1000) |
| NPM1 | Abcam | ab10530 | IF (1:500) |
| Alexa Fluor 488 Rb | Thermo Fisher | A-11008 | IF (1:500) |
| Alexa Fluor 647 Ms | Thermo Fisher | A-21235 | IF (1:500) |

| Kit | Source | Catalogue No |
|---|---|---|
| Click-iT® RNA Imaging Kit | Invitrogen | C10329, C10330 |
| DC Protein Assay Kit II | Bio-Rad | 5000112 |
| Thermo Scientific™ SuperSignal™ West Dura Extended Duration Substrate | Thermo Scientific | 10220294 |
| NEBridge® Golden Gate Assembly Kit (BsmBI-v2) | New England Biolabs (NEB) | E1601 |
| QIAprep Spin Miniprep Kit | Qiagen | 27104 |

| Chemical/Reagent | Company | Catalogue No | Solvent |
|---|---|---|---|
| Actinomycin D | Sigma Aldrich | A1410-10MG | DMSO |
| BMH21 | Sigma Aldrich | SML1183-5MG | DMSO |
| DMSO | Sigma Aldrich | D8418-250ML | - |
| Doxycycline | Sigma Aldrich | C8895 | dH2O |
| 3-Indoleacetic acid (Auxin) | Sigma Aldrich | I3750-5G-A | dH2O |
| Rapamycin | Sigma Aldrich | 37094 | DMSO |
| Hoechst 33342 | Sigma Aldrich | 14433 | DMSO |
| Polybrene | Sigma Aldrich | TR-1003-G | DMSO |
| Blasticidin | Thermo Fisher | R21001 | - |
| Geneticin (G418 Sulfate) | Thermo Fisher | 10131027 | - |
| Puromycin | Jena Bioscience | NU-931-05 | |
| Carbenicillin | Fisher Scientific | 11480952 | |
| Formaldehyde | Sigma Aldrich | F8775-500ML | |
| RIPA buffer | Thermo Fisher | PI-89901 | |
| Trypsin-EDTA | Thermo Fisher | 15400054 | |
| NuPAGE LDS Sample Buffer | Invitrogen | NP0007 | |
| NuPAGE Reducing Agent | Invitrogen | NP0009 | |
| T4 Ligation buffer | Roge | 11243292001 | |
| T4PNK | New England Biolabs (NEB) | M0201S | |
| Triton X-100 | Sigma Aldrich | X100 | |

| Target of sgRNA* | Primer Sequence Forward | Primer Sequence Reverse |
|---|---|---|
| RPA3 #1 | CACCGTACGGGTTCCATCAACTCGA | AAACTCGAGTTGATGGAACCCGTAC |
| RPA3 #2 | CACCGTAGCTTGATGAAGAAATCTC | AAACGAGATTTCTTCATCAAGCTAC |
| TP53 #1 | CACCGACTTCCTGAAAACAACGTTC | AAACGAACGTTGTTTTCAGGAAGTC |
| TP53 #2 | CACCGCTTACCAGAACGTTGTTTTC | AAACGAAAACAACGTTCTGGTAAGC |
| STAG2 #1 | CACCGCTTCAGTCGTAGAGATCCAG | AAACCTGGATCTCTACGACTGAAGC |
| STAG2 #2 | CACCGAGTCCCACATGCTATCCACA | AAACTGTGGATAGCATGTGGGACTC |
| TADA3 #1 | CACCGGGATCCGTGGCACGTCGATA | AAACTATCGACGTGCCACGGATCCC |
| TADA3 #2 | CACCGTCAGTAACTCCTCAAGTGTG | AAACCACACTTGAGGAGTTACTGAC |
| SUPT20H #1 | CACCGATTCATACCTTCCACTTGAT | AAACATCAAGTGGAAGGTATGAATC |
| SUPT20H #2 | CACCGTGTATACTTACCTTAACTTC | AAACGAAGTTAAGGTAAGTATACAC |
| SUPT20H #3 | CACCGCCCGACAGAGACCTCCTAAA | AAACTTTAGGAGGTCTCTGTCGGGC |
| TSC1 #1 | CACCGCGAGATAGACTTCCGCCACG | AAACCGTGGCGGAAGTCTATCTCGC |
| TSC1 #2 | CACCGGAGCATGTGCGAATTCATC | AAACGATGAATTCGCACATGCTCC |
| TSC1 #3 | CACCGACCTTCGAGGGTCCAGTTCA | AAACTGAACTGGACCCTCGAAGGTC |
| TSC2 #1 | CACCGTGGCCTCAACAATCGCATC | AAACGATGCGATTGTTGAGGCCAC |
| TSC2 #2 | CACCGAGCACGCAGTGGAAGCACTC | AAACGAGTGCTTCCACTGCGTGCTC |
| TSC2 #3 | CACCGTCTGCTGAAGGCCATCGTGC | AAACGCACGATGGCCTTCAGCAGAC |
| PTEN #1 | CACCGTTATCCAAACATTATTGCTA | AAACTAGCAATAATGTTTGGATAAC |
| PTEN #2 | CACCGCCTACCTCTGCAATTAAATT | AAACAATTTAATTGCAGAGGTAGGC |
| PTEN #3 | CACCGACCGCCAAATTTAATTGCAG | AAACCTGCAATTAAATTTGGCGGTC |

*Gene sequences taken from Ophir Shalem et al, 2014.
